# MAJIQ-SPEL: Web-Tool to interrogate classical and complex splicing variations from RNA-Seq data

**DOI:** 10.1101/136077

**Authors:** Christopher J. Green, Matthew R. Gazzara, Yoseph Barash

**Author notes:** These authors contributed equally to this work (Christopher J. Green, Matthew R. Gazzara).

## Abstract

Analysis of RNA sequencing (RNA-Seq) data have highlighted the fact that most genes undergo alternative splicing (AS) and that these patterns are tightly regulated. Many of these events are complex, resulting in numerous possible isoforms that quickly become difficult to visualize, interpret, and experimentally validate. To address these challenges, We developed MAJIQ-SPEL, a web-tool that takes as input local splicing variations (LSVs) quantified from RNA-Seq data and provides users with visualization and quantification of gene isoforms associated with those. Importantly, MAJIQ-SPEL is able to handle both classical (binary) and complex (non-binary) splicing variations. Using a matching primer design algorithm it also suggests users possible primers for experimental validation by RT-PCR and displays those, along with the matching protein domains affected by the LSV, on UCSC Genome Browser for further downstream analysis.

**Availability:** Program and code will be available at http://majiq.biociphers.org/majiq-spel

## 1 Introduction

Advances in RNA-Seq technology have led to improved detection and quantification of splicing variations through the use of short reads that span across spliced junctions. Most commonly used AS analysis tools focus exclusively on classical, binary AS events (e.g. cassette exon, alternative 5′ or 3′ splice sites, intron retention, etc.). Recently, we formulated local splicing variations (LSVs) that capture both classical as well as complex splicing patterns (i.e. involving three or more junctions). Surprisingly, we found that over 30% of splicing variations in extensive human and mouse RNA-Seq experiments we interrogated are complex [4].

The pervasiveness of complex splicing variations suggests that accurate interpretation of the underlying isoforms is crucial for experimentally interrogating and understanding the consequences of these splicing changes. We therefore developed MAJIQ and VOILA (http://majiq.biociphers.org) to define, quantify, and visualize such local splicing variations [4]. However, no current tool offers a user-friendly interface to connect LSVs, whether simple or complex, to the underlying known gene isoform and affected protein domains. Also, there is a clear need for automated design and visualization of potential primers that flank an LSV for experimental validation via RT-PCR, the gold standard in the field.

## 2 Results

We developed the web-tool MAJIQ-SPEL (MAJIQ for Sampling Primers and Evaluating LSVs) to aid in the visualization, interpretation, and experimental validation of both classical and complex splicing variations. MAJIQ-SPEL (or SPEL for short) is implemented on a Galaxy web server [1], taking as input the output of VOILA [4]. Specifically, users can now click a button to copy a splice graph and LSV quantification of interest, then paste it into SPEL’s Galaxy input form.

MAJIQ-SPEL output contains several components, which we highlight in Figure 1 using a complex LSV within *Clta* generated comparing RNA-Seq from mouse cerebellum and adrenal gland [5]. First, colorized representations of the LSV are displayed with junction spanning read counts for each junction quantified directly in the LSV (colored arcs) as well as those that occur within the boundaries of the event, but are not directly quantified (dashed grey arcs) (Figure 1A). This allows for quick interpretation of which paths are commonly utilized in each sample. Second, SPEL produces a table of putative forward and reverse primers for the 5′- and 3′-most exons within the LSV (Figure 1B). The primers are optimized for validating the given LSV via low-cycle RT-PCR and their design can be adjusted using user specified parameters such as length, GC content, melting temperature (T_m_) formula and range. The algorithm implemented in SPEL for the primer design is based on and the experimental protocols and primer design factors described in [3]. For ease of use, the primer table is searchable and key summary information for each primer can also be displayed.

**Figure 1:**
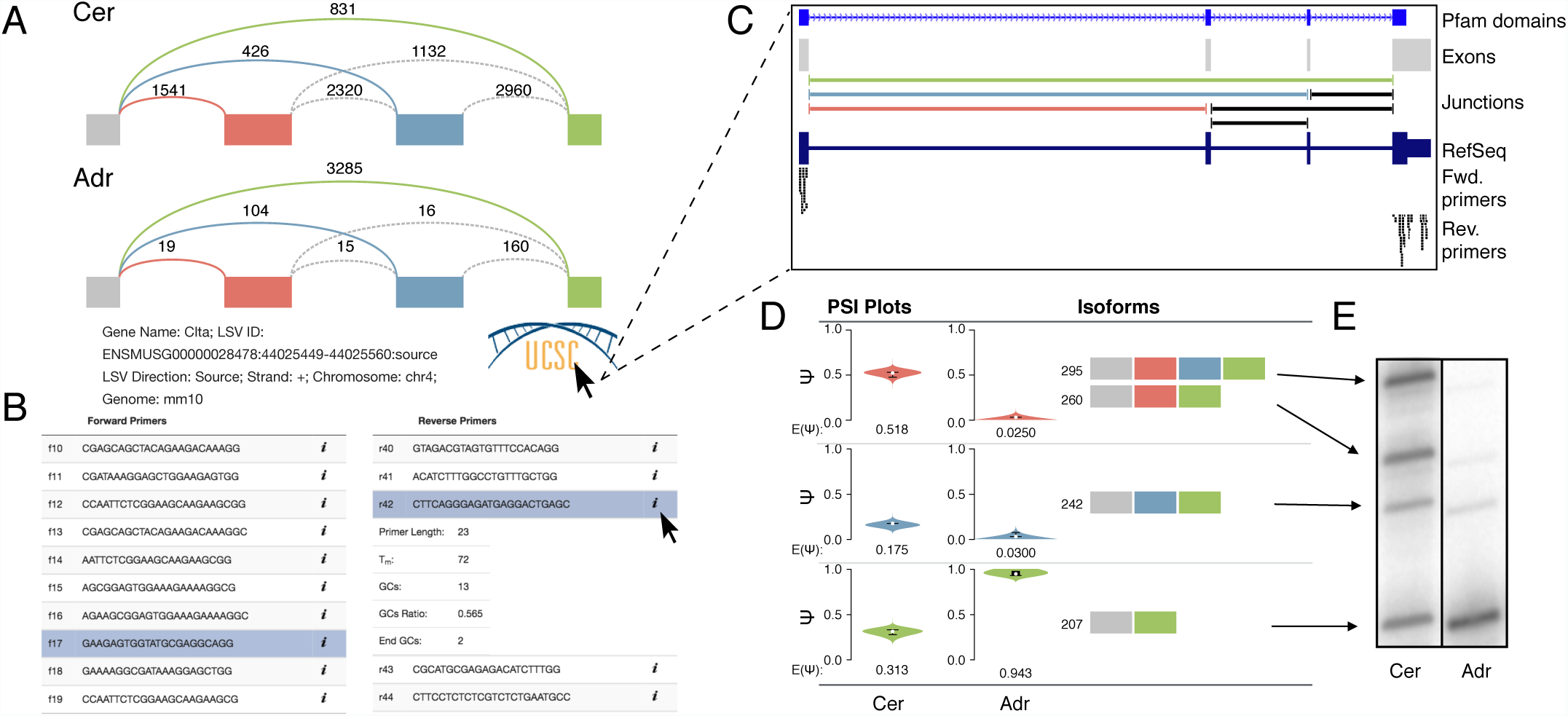
**(A)** Splice graph representation of LSV within *Clta* from mouse cerebellum (top) and adrenal gland (bottom) with reads detected from RNA-Seq data displayed above each junction. Junctions quantified directly within the LSV are colored. **(B)** Primer table that suggests possible forward (left) and reverse (right) primers. Additional information for each primer can be displayed by clicking the ”i”. **(C)** UCSC Genome Browser snapshot with custom tracks produced by MAJIQ-SPEL as labeled. **(D)** Isoform table that displays PSI quantifications (left) and possible isoforms (right). Nucleotide sizes correspond to products produced using the selected primers from (B). **(E)** Representative RT-PCR validation of predicted product sizes and quantification using the primers selected in (B) on total RNA from mouse cerebellum (left) and adrenal gland (right).

MAJIQ-SPEL also offers UCSC Genome Browser connectivity. Clicking the browser’s logo brings up custom tracks that display the exons, junctions, and locations of putative primers to aid in selection of primers for validation (Figure 1C). These tracks also include annotated protein domains (Pfam [2]), which can aid in examining the functions of alternative isoforms produced.

Finally, MAJIQ-SPEL traverses all possible paths within the splice graph to generate the isoform table that links the MAJIQ PSI (percent selected index, Ψ) quantification in each condition to the associated isoform(s) (Figure 1D). Importantly, once the user selects a forward and reverse primer pair, this table updates to display the expected product size for each isoform for validation. RT-PCR performed using primers generated by MAJIQ-SPEL demonstrates both accurate prediction of all four product sizes and quantification for both cerebellum and adrenal gland (Figure 1E).

Beyond handling classic or complex splicing variations, MAJIQ-SPEL also offers researchers fast and accurate primer design for de novo splicing variations not in the annotated transcriptome. In such cases experimental validation is crucial. Such a case is shown in an event in *Fubp3* (Figure S1). Since this LSV involves novel exon skipping it will likely not be captured in other tools for splicing quantification and visualization packages that rely only on the annotation database.

The *Fubp3* and *Clta* splicing variations shown here also highlight how MAJIQ-SPEL can aid in functional analysis of LSVs. The combined UCSC Genome Browser tracks show the alternative exons overlap annotated protein domains, suggesting a functional effect. The cassette exon in *Fubp3* is not a multiple of three, suggesting a frameshift and the Browser tracks revealed that skipping inserts a premature termination codon (PTC). Future extensions of this work will aim to further integrate these and other functional analyses into MAJIQ-SPEL.

## Acknowledgments

Special thanks to Jorge Vaquero-Garcia, Anupama Jha, and Kristen W. Lynch for support and advice throughout this project.

---

**Figure S1:**
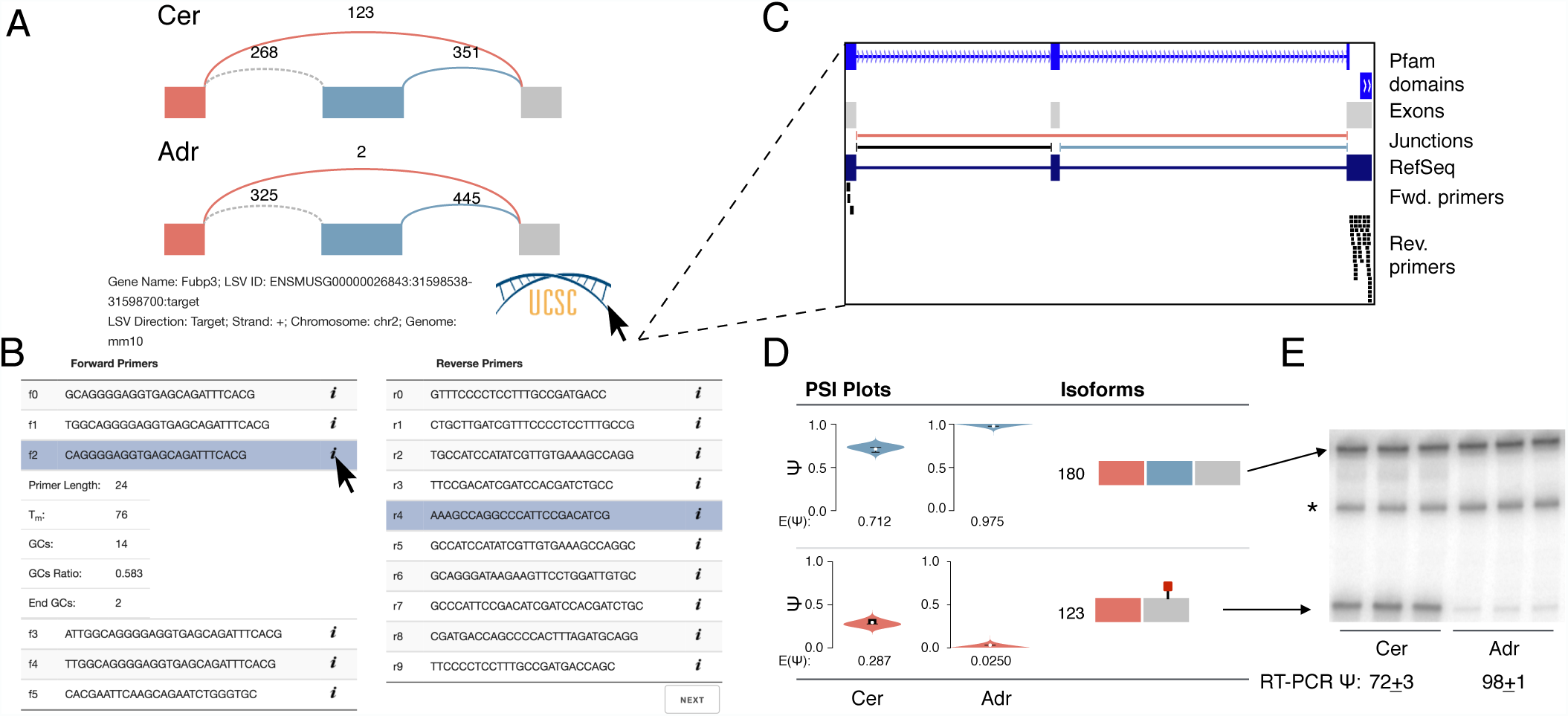
**(A)** Splice graph representation of LSV involving novel exon skipping event within *Fubp3* from mouse cerebellum (top) and adrenal gland (bottom) with reads detected from RNA-Seq data displayed above each junction. Junctions quantified directly within the LSV are colored. **(B)** Primer table that suggests possible forward (left) and reverse (right) primers. Additional information for each primer can be displayed by clicking the ”i”. **(C)** UCSC Genome Browser snapshot with custom tracks produced by MAJIQ-SPEL as labeled. Pfam domains show this alternative exon overlaps with the first annotated KH domain, suggesting skipping could affect RNA binding of Fubp3. **(D)** Isoform table that displays PSI (Ψ) quantifications (left) and possible isoforms (right). Nucleotide sizes correspond to products produced using the selected primers from (B). Red stop sign indicates premature termination codon (PTC) introduction upon exon skipping that may induce nonsense mediated decay (NMD). **(E)** RT-PCR validation of predicted product sizes and quantification using the primers selected in (B) on total RNA from mouse cerebellum (left) and adrenal gland (right). Average inclusion of the cassette exon by RT-PCR with standard deviation is given. Asteriks corresponds to a background band that does not run true to size.

